# Scaling and Generalization of Discrete Diffusion Models for Tumor Phylogenies

**DOI:** 10.64898/2026.03.23.713822

**Authors:** Siddharth Sabata, Russell Schwartz

## Abstract

Tumor phylogenies — rooted trees encoding clonal ancestry and mutation acquisition — are central to understanding cancer evolution, yet generating realistic phylogenies remains challenging. We investigate whether discrete graph diffusion can learn the structural constraints of tumor phylogenies directly from data. Working with approximately 12,500 synthetic phylogenies across twelve evolutionary regimes, we train graph transformer models that denoise typed graphs through a learned reverse diffusion process. Scaling experiments reveal a non-monotonic capacity–performance relationship: a mid-scale model achieves high structural validity and close distributional match to held-out data, while a deeper model fails under fixed optimization hyperparameters. Low-data cross-regime experiments show that diverse training produces more transferable representations than single-regime specialization. These results establish that phylogenetic structural constraints can be learned implicitly through unconditional discrete diffusion, suggesting a viable path toward generative models of tumor evolution.

## 1 Introduction

Tumor phylogenies—trees encoding the clonal ancestry and mutational history of a tumor—are central to understanding treatment resistance, predicting disease trajectories, and identifying therapeutic targets [1, 7]. Deep generative models have advanced other areas of structural biology, including protein structure prediction [12] and de novo protein design [20], yet tumor phylogenies remain largely outside the scope of these methods. Recent work has applied deep generative models to species phylogenetics—using VAEs [22], GFlowNets [24], and language-model-guided methods [4] — but these target molecular sequence phylogenies and have not been adapted to the distinct structural constraints of tumor evolution. Here, we ask the question of whether discrete graph diffusion can learn structural rules governing tumor phylogenies from data alone?

Tumor phylogenies are typed clone–mutation graphs subject to strict structural constraints: acyclicity, a single root, categorically typed nodes (root, clone, mutation) and edges (clone ancestry, mutation assignment), and variable graph sizes across patients. Moreover, evolutionary dynamics vary qualitatively — from slow clonal sweeps to rapid branching evolution — so a useful generative model must capture structural regularities that hold across diverse tumor types. Inference methods such as PhyloWGS [3], Canopy [11], and SCITE [10] reconstruct trees from sequencing data through inference-time optimization (MCMC or tree enumeration), but face scalability limitations as clone counts grow. These limitations motivate amortized approaches that shift computational effort from inference to training time. Discrete diffusion for graphs [2, 19] provides a natural framework for generating typed graph structures capturing the essential features of phylogenies.

We present **DiPhy** (Discrete diffusion for Phylogenies), which adapts discrete graph diffusion to tumor phylogeny generation. We construct a synthetic dataset of ∼ 12,500 tumor phylogenies spanning 12 biologically distinct evolutionary regimes using the SISTEM simulator [21] with Latin hypercube sampling [14] for systematic parameter coverage. For the present work, we focus on the problem of unconditional generation — learning to produce phylogenies without conditioning on observed data from a specific evolutionary trajectory — because it reveals what structural patterns diffusion can capture from training data alone and directly tests whether strict phylogenetic constraints (acyclicity, single root, typed node–edge consistency) emerge from distributional learning or require explicit enforcement. We evaluate DiPhy’s ability to learn to generate samples over a space of tumor phylogeny-like graphs through scaling ablations over model capacity and data size, and cross-regime generalization experiments.

Our main contributions are:

- **Representation**. A typed clone–mutation graph encoding for tumor phylogenies compatible with discrete graph diffusion, with categorical node types (root, clone, mutation) and edge types (clone ancestry, mutation assignment).
- **Data**. A synthetic benchmark dataset of ∼ 12,500 phylogenies spanning 12 SISTEM regimes with systematic parameter coverage via Latin hypercube sampling.
- **Empirical characterization**. Scaling ablations over model depth × data size; cross-regime generalization experiments; and analysis of validity versus distributional fidelity, which exhibit partial dissociation with distinct scaling behavior.
- **Release**. Code and datasets are publicly available on GitHub.

## 2 Related Work

The specific task undertaken in the present work, of learning a diffusion model of tumor phylogeny-like graph structure, is to our knowledge, novel. However, a number of related directions have been explored in the literature.

There is an extensive literature now on phylogeny methods for inferring clonal lineages from tumor phylogeny data. See, for example, [17] for an overview of popular approaches to the problem. Traditional methods formulate phylogeny reconstruction as probabilistic inference. PhyloWGS [3] infers clonal phylogenies from bulk sequencing via MCMC; Canopy [11] uses EM-style optimization with tree enumeration; SCITE [10] models mutation accumulation from single-cell data. These methods face scalability limitations as clone counts grow, motivating amortized learning approaches. No prior work applies discrete graph diffusion to tumor phylogenies or studies how training regime diversity affects structural validity and distributional fidelity in this domain.

Our work specifically builds on recent work on discrete diffusion models for graphs. D3PM [2] introduced discrete diffusion by replacing Gaussian transitions with Markov chains over categorical states. DiGress [19] extended this to graphs, treating node and edge types as independent categorical variables with marginal-based transitions and graph transformer architectures. DiGress achieves high validity on molecular benchmarks without explicit constraint enforcement. Whether implicit learning transfers to domains with strict hierarchical constraints—like phylogenies requiring global acyclicity and typed consistency—is the question we address.

Several recent works have applied deep generative models to phylogenetic tree generation. PhyloVAE [22] uses a variational autoencoder with autoregressive tree generation for multivariate phylogenetic data. PhyloGFN [24] frames Bayesian phylogenetic inference as a GFlowNet problem, sampling trees proportional to posterior probability. PhyloGen [4] combines language model embeddings with graph structure generation. phyloGAN [18] applies GANs to small phylogenetic trees but reports training instability limiting scalability. All of these target molecular sequence phylogenies rather than tumor phylogenies.

Other forms of neural network model have also recently been explored for phylogeny inference tasks. Phyloformer [15] uses transformers for fast phylogenetic reconstruction matching maximum likelihood accuracy. NeuralNJ [23] provides a differentiable neighbor-joining algorithm with learned distance corrections. Phyla [6] trains a hybrid state-space and transformer model with phylogenetic supervision across protein families. These inference methods accelerate tree reconstruction from observed sequences; DiPhy instead generates phylogenies unconditionally and targets tumor clone trees with typed nodes and edges.

## 3 Data and Representation

This section defines the typed-graph encoding used for tumor phylogenies and describes the synthetic dataset on which DiPhy is trained.

### 3.1 Tumor Phylogenies as Typed Graphs

A **tumor phylogeny** is a rooted tree where each internal node represents a clone (a genetically distinct subpopulation), edges encode direct ancestral relationships, and the root represents normal cells. Clones accumulate mutations; descendant clones inherit all ancestral mutations plus new ones, producing a nested structure.

Formally, a clone tree *T* = (*V*_*C*_, *E*_*C*_, *r*) consists of clone nodes *V*_*C*_ = {*r, c*_1_, …, *c*_*k*_} with root *r*, directed edges *E*_*C*_ ⊆ *V*_*C*_ *× V*_*C*_, and a mutation assignment *M* : *V*_*C*_ → 2^ℳ^ satisfying the inheritance constraint: if *c*^*′*^ descends from *c*, then *M* (*c*) ⊆ *M* (*c*^*′*^).

#### Typed-Graph Encoding

Clone trees with variable-sized mutation sets are difficult to process with graph neural networks. We adopt an **unrolled typed-graph encoding** that makes mutations explicit as nodes (Figure 1). Given a clone tree *T* with mutation assignment *M*, we construct a typed graph *G* = (*X, E*) with:

- **Node types** *X* ∈ { 0, 1, 2 }^*n*^: type 0 = root (single normal-cell node), type 1 = clone, type 2 = mutation (one per mutation event).
- **Edge types** *E* ∈ { 0, 1, 2 }^*n×n*^ (symmetric): type 0 = no edge, type 1 = clone edge (root–clone or clone–clone ancestry), type 2 = mutation edge (clone–mutation assignment).

**Figure 1:**
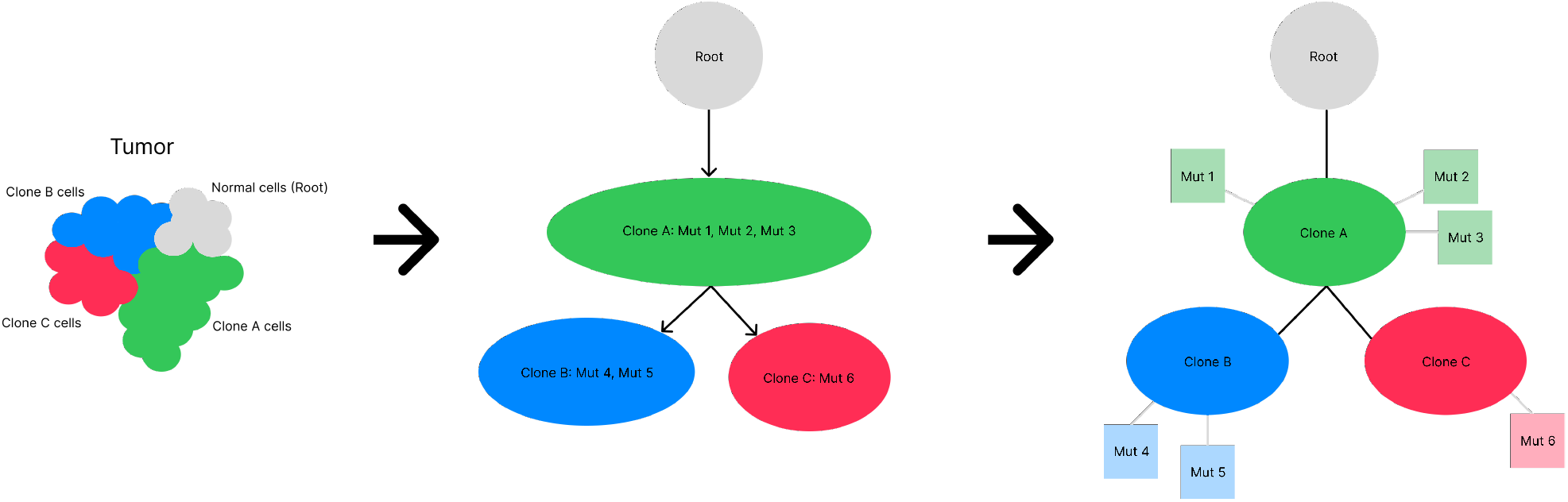
Graph encoding from simulator output to model input. A tumor phylogeny with nested mutations (left) is converted to an unrolled typed graph (right). The root node (gray) represents normal cells. Clone nodes (colored, bold) form a tree backbone. Each mutation is extracted as an individual node (small circles) connected to its defining clone via mutation edges (light gray). Clone-clone edges (black) become undirected in the final encoding.

**Figure 2:**
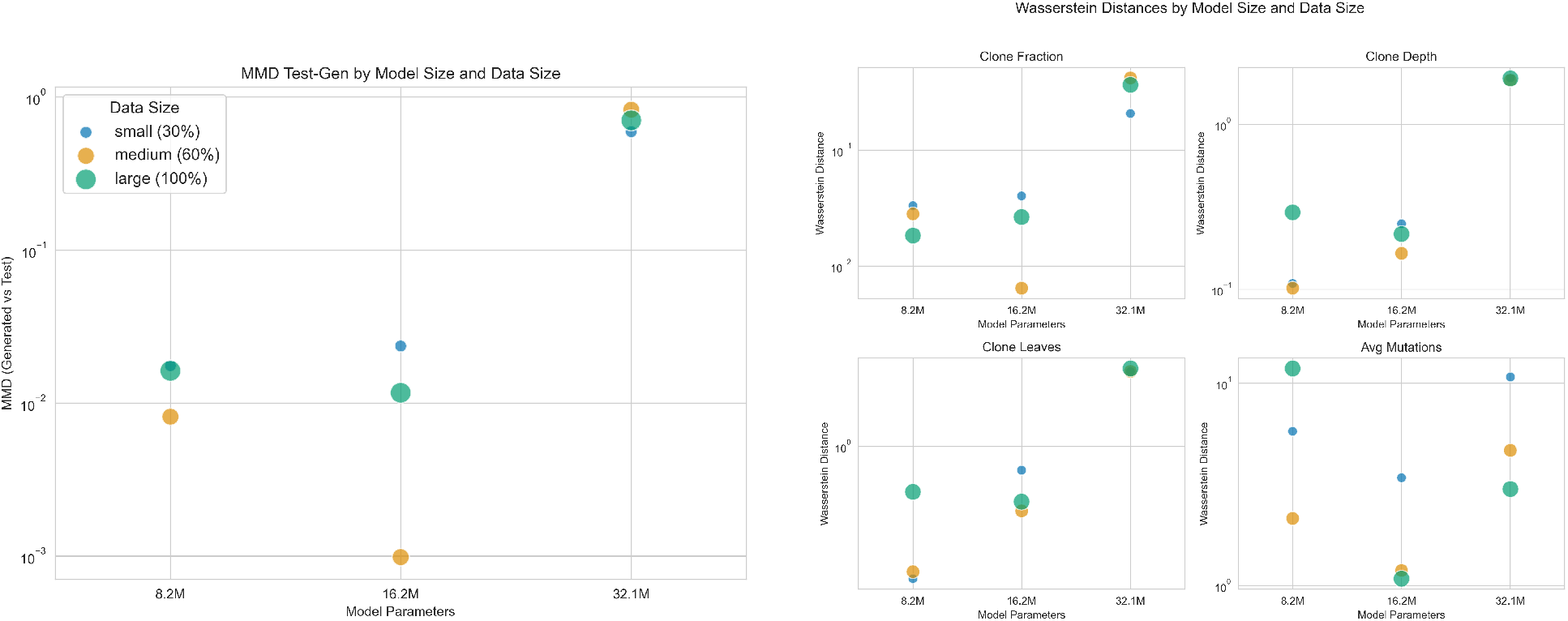
Distributional fidelity across model and data size. (Left) MMD^2^ between generated and held-out test distributions as a function of training set size. (Right) Per-feature Wasserstein distances across scaling configurations, showing feature-level distributional agreement.

### 3.2 SISTEM Dataset

We generate a synthetic training data set of trees meant to capture the essential features of tumor phylogenies using SISTEM [21], which simulates stochastic cell division, mutation acquisition, and selection from a single founding cell. SISTEM produces ground-truth clone trees with mutation assignments; we use only the tree topologies produced by SISTEM, disregarding the synthetic sequencing data it also generates associated with each tree.

#### Regime Design

Our intention is to train a graph diffusion model that is capable of generating a diverse array of tumor phylogeny-like tree topologies. To avoid learning simulator-specific artifacts from a single configuration, we define twelve evolutionary regimes spanning a variety of biologically plausible tumor dynamics (Table 4, Appendix):

- **Single-site primary tumors (R01–R05):** varying selection strength, Copy Number Alteration (CNA) types, and chromosomal instability.
- **Three-site metastatic tumors (R06–R07):** introducing inter-site migration with shared or distance-based fitness landscapes.
- **Five-to-seven-site metastatic tumors (R08–R11):** complex multi-site spread with high migration, genotype-dependent migration, and organotropism.
- **Small trees / early detection (R12):** low-burden tumors producing structurally simple trees.

Within each regime, continuous parameters (mutation rates, fitness coefficients, detection thresholds) are sampled via Latin Hypercube Sampling [14] for systematic coverage of the local parameter space.

The final dataset comprises 12,581 phylogenies after filtering. Graphs range from 5 to 200 nodes, with clone fraction spanning 0.15–0.45 across regimes, maximum clone depth from 2 to 12, and branching factor typically 1.5–2.5. To control memory costs from *O*(*n*^2^) edge tensors, graphs exceeding 200 nodes are excluded, primarily affecting metastatic regimes R08–R11 with large trees; regime R08 retains only 64 of 1500 planned samples due to the low success rate of producing samples that result in stable multi-site growth. A small number of structurally malformed simulator outputs (disconnected components, missing root) are also removed. These filters bias the dataset toward smaller phylogenies.

Each phylogeny is encoded as a typed graph following the procedure above and stored as a dictionary containing *X, E*, and human-readable labels *L*. Only *X* and *E* are used for training. Data is split 80/10/10 (train/validation/test) with a fixed random seed.

## 4 Method: Discrete Diffusion on Typed Phylogeny Graphs

DiPhy adapts the DiGress discrete graph diffusion framework [19] to unconditional tumor phylogeny generation. We describe the diffusion process, architecture, and training configuration, focusing on aspects that differ from or extend DiGress. See Appendix **??** for a full correspondence table.

### 4.1 Discrete Graph Diffusion

Given a clean graph *G*_0_ = (*X*_0_, *E*_0_) with one-hot node types *X*_0_ ∈ { 0, 1 }^*n×*3^ and edge types *E*_0_ ∈ { 0, 1 }^*n×n×*3^, the forward process corrupts categories through Markov transition matrices over *T* = 1000 timesteps:

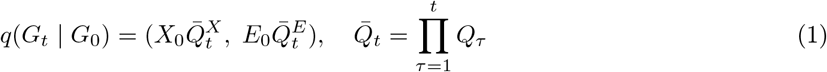

#### Marginal Transitions

We use marginal-preserving transitions rather than uniform transitions:

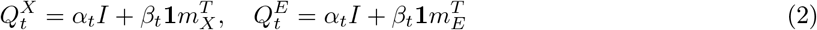

where *m*_*X*_, *m*_*E*_ are empirical marginals from the training set and *α*_*t*_ + *β*_*t*_ = 1. This is important for phylogenies: edge type distributions are highly imbalanced (*>*95% no-edge entries), so uniform transitions would destroy sparsity structure early in the forward process.

#### Reverse Process

A graph transformer *ϕ*_*θ*_ predicts the clean graph *Ĝ*_0_ from noisy input *G*_*t*_ and time step *t*. Generation proceeds by sampling from the posterior:

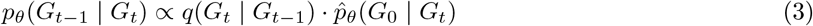

### 4.2 Architecture

DiPhy uses a graph transformer blocks [5]that jointly process node, edge, and global representations through self-attention. Edge features are modulated by global features, and node features by both edge and global features, via Feature-wise Linear Modulation (FiLM) [16]. Edges are kept symmetric (*e*_*ij*_ = *e*_*ji*_). The normalized diffusion time step *t/T* is concatenated to the global feature vector. Separate MLP heads project final representations to categorical distributions over node and edge types.

### 4.3 Model Configurations

We evaluate three model sizes (Table 1), scaling capacity through depth only while fixing hidden dimensions (*d*_*X*_ = 256, *d*_*E*_ = *d*_*y*_ = 64, 8 heads). All configurations share identical hyperparameters except batch size, which is halved per depth doubling (28, 14, 7) to fit in GPU memory; the implications for the 32.1M model are analyzed in Section 7.1.

**Table 1:**
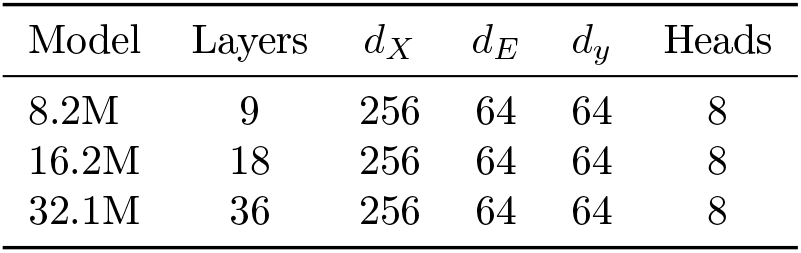
Model configurations used in scaling experiments.

| Model | Layers | $d_X$ | $d_E$ | $d_y$ | Heads |
| --- | --- | --- | --- | --- | --- |
| 8.2M | 9 | 256 | 64 | 64 | 8 |
| 16.2M | 18 | 256 | 64 | 64 | 8 |
| 32.1M | 36 | 256 | 64 | 64 | 8 |

### 4.4 Training

We train with cross-entropy loss on node and edge predictions:

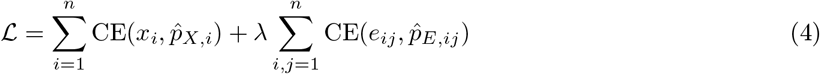

with *λ* = 5 to upweight edge predictions, reflecting the *O*(*n*^2^) edge space with highly imbalanced type distributions. Global features are disabled (*λ*_*y*_ = 0); graph-level information is learned implicitly through pooling.

All models are trained for 500 epochs with AdamW with AMSGrad (learning rate 2 × 10^−4^, weight decay 10^−12^), batch size 28 for the 9-layer model, and no learning rate schedule. The best checkpoint is selected by validation NLL. Training uses NVIDIA H100 80GB GPUs. The 8.2M model trains in ∼ 24 hours, the 16.2M in ∼ 36 hours on a single H100 using the full dataset. Memory management details (sharding, on-the-fly encoding) are described in Appendix B.

## 5 Experiments

We conduct two sets of experiments: scaling ablations that vary model and data size, and a low-data regime-coverage experiment that probes generalization under distribution shift.

### 5.1 Scaling Ablations

We train models of three sizes (8.2M, 16.2M, and 32.1M parameters) on three data fractions (30%, 60%, and 100% of the full dataset of 12,581 graphs), yielding nine configurations. Data fractions are created by stratified random subsampling that preserves regime proportions, using a fixed random seed.

### 5.2 Low-Data Regime-Coverage Experiment

This experiment probes whether models learn transferable phylogenetic structure or memorize regime-specific patterns. We use 700 training graphs to force a representation-sharing regime where memorization is less feasible, highlighting transfer effects. Specifically, we train 8.2M parameter models on 700 graphs under three regime-coverage conditions. The **Regular** condition uses uniform sampling across all 12 regimes ( ∼ 58 graphs per regime). The **R1 Only** condition draws all 700 graphs from regime R01 (single-site, near-neutral evolution). The **No R1** condition uses uniform sampling from regimes R02–R12, holding out R01 entirely.

### 5.3 Evaluation Metrics

All experiments are evaluated on 1,000 generated graphs using the following metrics.

#### 5.3.1 Structural Validity

A generated graph *G* = (*X, E*) is **phylogenetically valid** if it satisfies four constraints:

**Constraint 1: Acyclicity**. The clone subgraph *G*_*C*_ = (*V*_clone_, *E*_clone_) must contain no cycles:

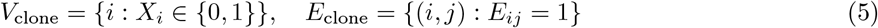

**Constraint 2: Single Root with Degree One**.

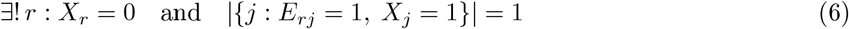

**Constraint 3: Clone Edge Validity**. Clone edges connect only root or clone nodes:

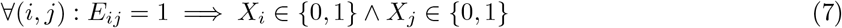

**Constraint 4: Mutation Edge Validity**. Mutation edges connect exactly one clone to one mutation node:

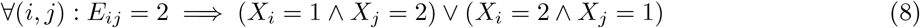

Overall validity requires all four constraints. We report both aggregate and per-constraint pass rates.

#### 5.3.2 Distributional Metrics

For each graph *G* = (*X, E*), we extract a feature vector **f** (*G*) ∈ ℝ^4^:

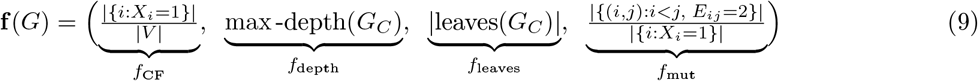

where *G*_*C*_ is the clone subtree (nodes with *X*_*i*_ ∈ {0, 1 } connected by clone edges *E*_*ij*_ = 1). The four components are clone fraction, maximum clone depth, number of clone leaves, and average mutations per clone.

We compute per-feature 1-Wasserstein distances between generated samples 𝒮_gen_ and held-out test samples 𝒮_test_:

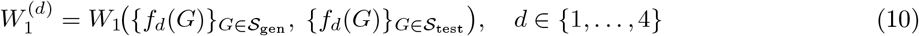

For a global summary, we compute the unbiased squared Maximum Mean Discrepancy [9] between generated and test distributions over the full feature vector. Let 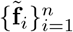 and 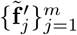 denote the z-score standardized features (zero mean, unit variance computed over all generated, training, and test samples) for the *n* generated and *m* test graphs, respectively. The estimator is:

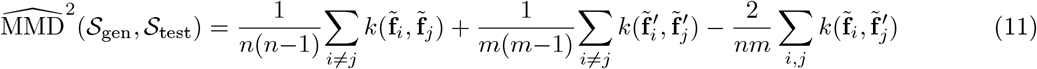

where *k*(**x, y**) = exp( −∥**x** −**y**∥ ^2^*/*2*σ*^2^ is the RBF kernel with bandwidth *σ* set to the median pairwise Euclidean distance over training and test features.

### 5.4 Implementation Details

All experiments use seed 0 for data splitting and model initialization. Models are trained for 500 epochs, and the best checkpoint by validation negative log likelihood (NLL) is used for evaluation. At inference time, we generate 1,000 graphs using 1,000 diffusion steps with no classifier-free guidance or sampling modifications.

## 6 Results

### 6.1 Scaling Behavior

Table 2 consolidates validity, distributional fidelity (MMD^2^), and representative per-feature Wasserstein distances across scaling configurations. Table 3 provides the full per-constraint validity breakdown.

**Table 2:**
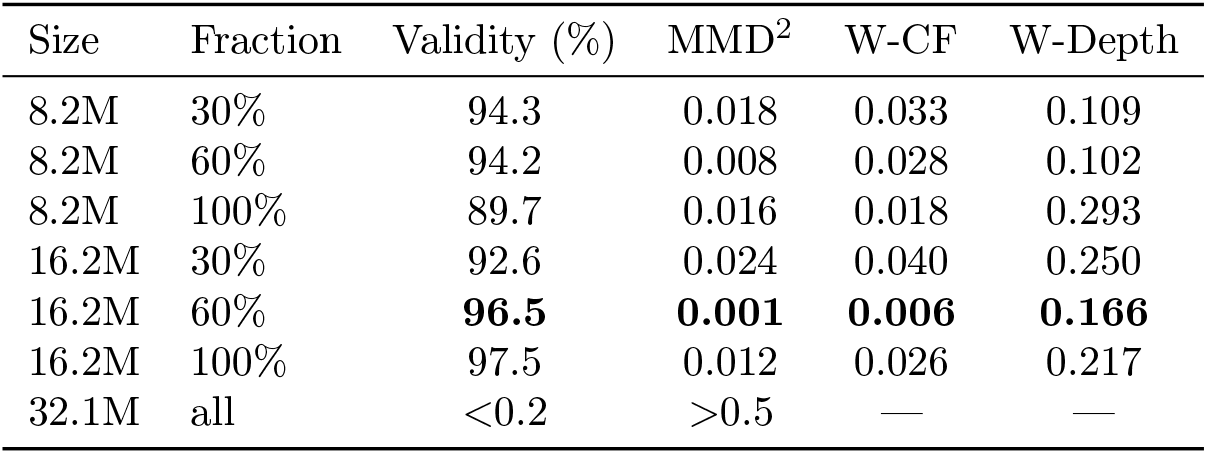
Consolidated scaling results. Validity (% of 1,000 generated graphs satisfying all four constraints), MMD^2^ between generated and test distributions, and 1-Wasserstein distances on clone fraction (W-CF) and tree depth (W-Depth). The 32.1M model diverged across all data fractions (Section 7.1). Bold indicates best per column among non-diverged models.

| Size | Fraction | Validity (%) | MMD<sup>2</sup> | W-CF | W-Depth |
| --- | --- | --- | --- | --- | --- |
| 8.2M | 30% | 94.3 | 0.018 | 0.033 | 0.109 |
| 8.2M | 60% | 94.2 | 0.008 | 0.028 | 0.102 |
| 8.2M | 100% | 89.7 | 0.016 | 0.018 | 0.293 |
| 16.2M | 30% | 92.6 | 0.024 | 0.040 | 0.250 |
| 16.2M | 60% | **96.5** | **0.001** | **0.006** | **0.166** |
| 16.2M | 100% | 97.5 | 0.012 | 0.026 | 0.217 |
| 32.1M | all | <0.2 | >0.5 | — | — |

**Table 3:** Per-constraint validity across scaling configurations. Percentage of 1,000 generated graphs satisfying each constraint. Bold indicates best per data fraction (excluding diverged 32.1M).

| Size | Fraction | Overall | No Cycle | Root Degree | Clone Edge | Mutation Edge |
| --- | --- | --- | --- | --- | --- | --- |
| 8.2M | 30% | 94.3 | 96.0 | 97.6 | 99.5 | 99.7 |
| 16.2M | 30% | 92.6 | 95.8 | 96.0 | 99.3 | 99.6 |
| 8.2M | 60% | 94.2 | 94.8 | 97.1 | 99.9 | 99.7 |
| 16.2M | 60% | **96.5** | **97.6** | **98.2** | **100.0** | **99.9** |
| 8.2M | 100% | 89.7 | 92.5 | 95.6 | 99.3 | 99.2 |
| 16.2M | 100% | **97.5** | **98.5** | **99.2** | 99.4 | **99.8** |

Scaling ablations reveal a non-monotonic relationship between model capacity and generative performance. The 16.2M model at 60% data achieved the best overall balance: 96.5% validity with MMD^2^ = 0.001. Increasing to 100% data raised validity to 97.5% but increased MMD^2^ to 0.012, consistent with mild overfitting to training-specific patterns. By contrast, the smaller 8.2M model maintained 89–94% validity across data scales but showed higher Wasserstein distances, indicating underfitting. Its validity decreased from 94.2% at 60% data to 89.7% at 100%, consistent with the full dataset introducing structural diversity that exceeds the smaller model’s capacity.

At the high end of the capacity range, the 32.1M model (36 layers) achieved near-zero validity across all data fractions. Training logs show immediate divergence at 30% and 60% data (best checkpoint at epoch 0) and late divergence at 100% data (epoch 149), consistent with optimization instability when scaling transformer depth 4 × without adapting the learning rate or adding stabilization (warmup, gradient clipping, Pre-LayerNorm); Section 7.1 provides a detailed analysis. Across all non-diverged configurations, edge type constraints were easiest to satisfy (clone and mutation edge validity *>*99%), followed by root degree (95–99%) and acyclicity (92–98%). This ordering reflects the locality of each constraint: edge-type rules depend on adjacent node pairs, while cycle-freedom requires global path reasoning.

### 6.2 Cross-Regime Generalization

We trained 8.2M models on 700 graphs under three conditions: “Regular” (uniform across all 12 regimes), “R1 Only” (regime 1 exclusively), and “No R1” (regimes 2–12, holding out regime 1). Full metrics appear in Table 6 (Appendix).

Single-regime training (R1 Only) achieved the highest validity (66.2%) but poor generalization: Figure 3 shows elevated MMD on non-R1 regimes. The Regular model achieved lower validity (40.9%) but more uniform error across regimes, revealing a trade-off between in-distribution validity and cross-regime generalization. This pattern extends to held-out regime transfer: the No R1 model (46.9% validity) partially generalized to the unseen R1 regime, with MMD on R1 (0.15) elevated relative to trained regimes (0.02–0.05) but lower than the R1 Only model’s error on non-R1 regimes (*>*0.2). Structure learned from diverse regimes appears to transfer more readily than structure from a single regime.

**Figure 3:**
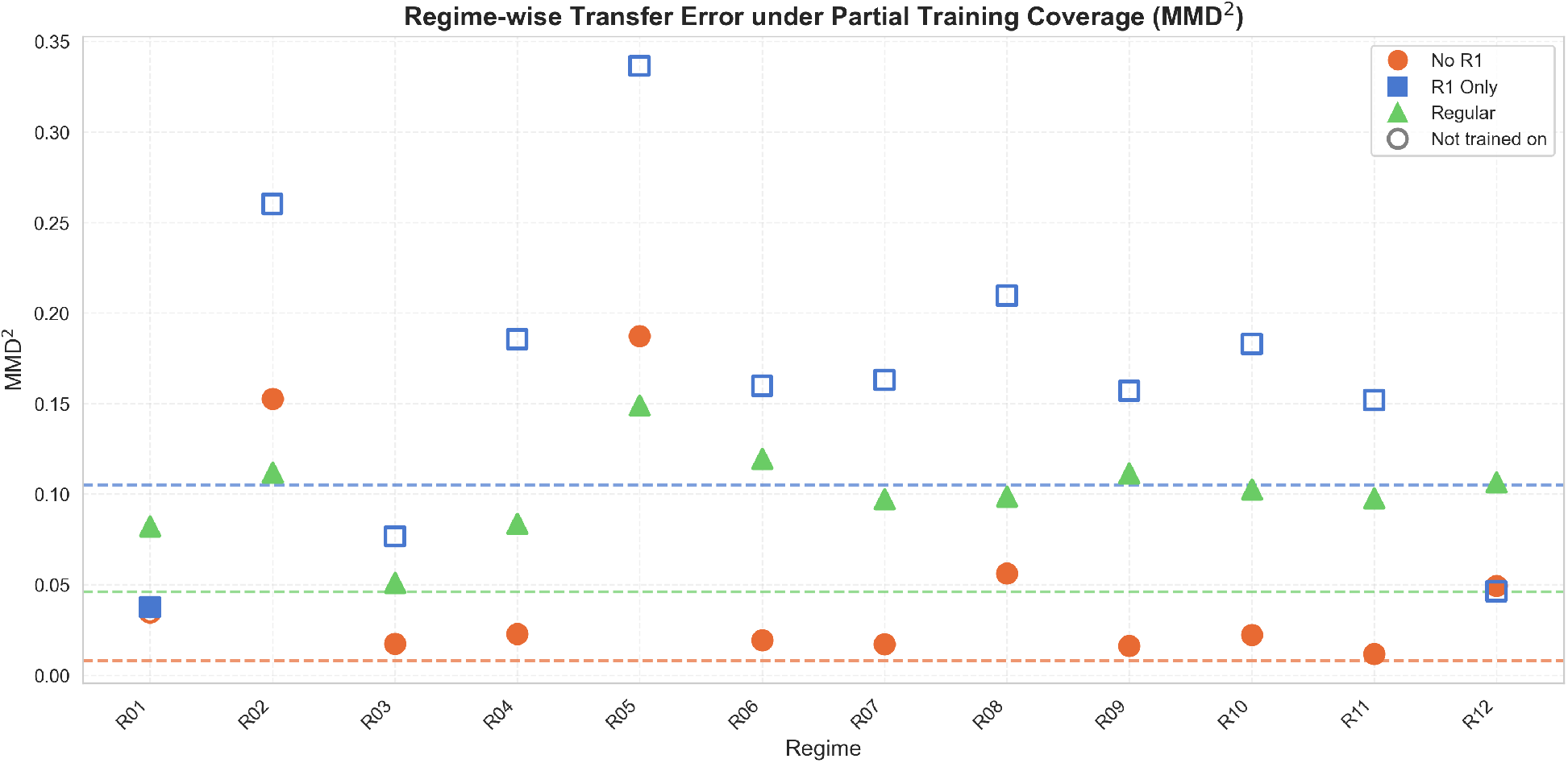
Cross-regime generalization. MMD^2^ between generated samples and each of the 12 test regimes for three training configurations: “Regular” (all 12 regimes, triangles), “R1 Only” (regime 1 only, squares), and “No R1” (regimes 2–12, circles). Hollow markers indicate regimes absent from training. Dashed lines show overall MMD^2^. The R1 Only model specializes but fails to transfer; the No R1 model partially generalizes to the held-out regime.

**Figure 4:**
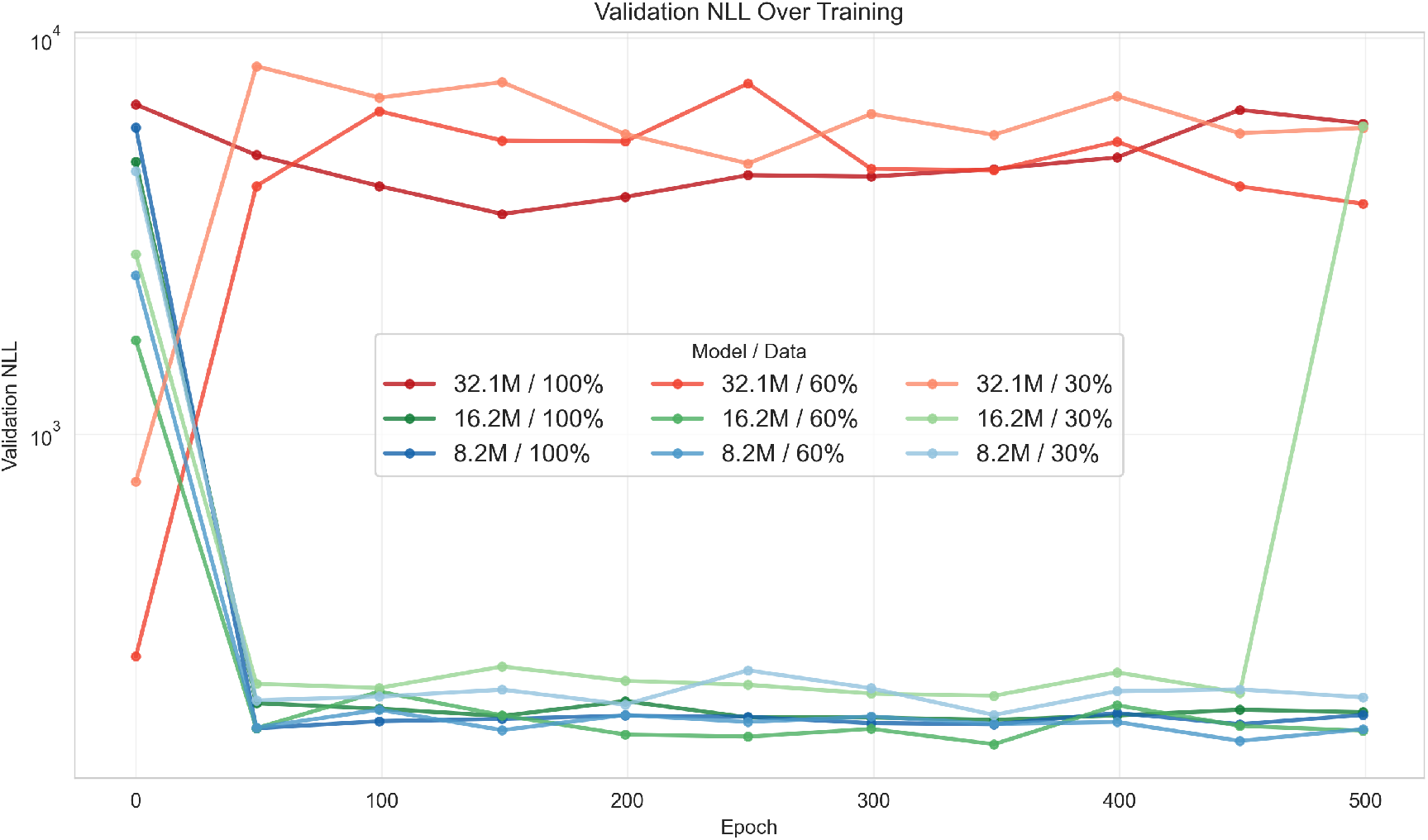
Validation loss trajectories across scaling configurations. Negative log-likelihood (NLL) on the validation set over 500 training epochs for all model size and data fraction combinations. The 8.2M and 16.2M models show stable convergence across data scales, with validation NLL decreasing to 165–220 and remaining stable. In contrast, all 32.1M configurations exhibit pathological behavior: the model trained on 30% data achieves its best validation loss at epoch 0 and diverges immediately; the 60% configuration shows similar immediate divergence; even with 100% data, the 32.1M model’s validation loss increases after epoch 149, indicating optimization instability (Section 7.1).

Figure 3 also shows systematic variation in per-regime difficulty. Metastatic regimes (R06–R10) showed higher error across all models, while single-site regimes (R01–R05) were easier to match, and R12 (small trees) was well-captured by all models. Generation difficulty correlates with phylogenetic complexity. This is consistent with the feature-space structure shown in Figure 5: R01 and R12 occupy nearby regions in the PCA projection, reflecting shared structural simplicity (shallow trees, low clone fraction). The R1 Only model’s ability to partially capture R12 suggests it learned a structural motif common to both regimes rather than regime-specific details.

**Figure 5:**
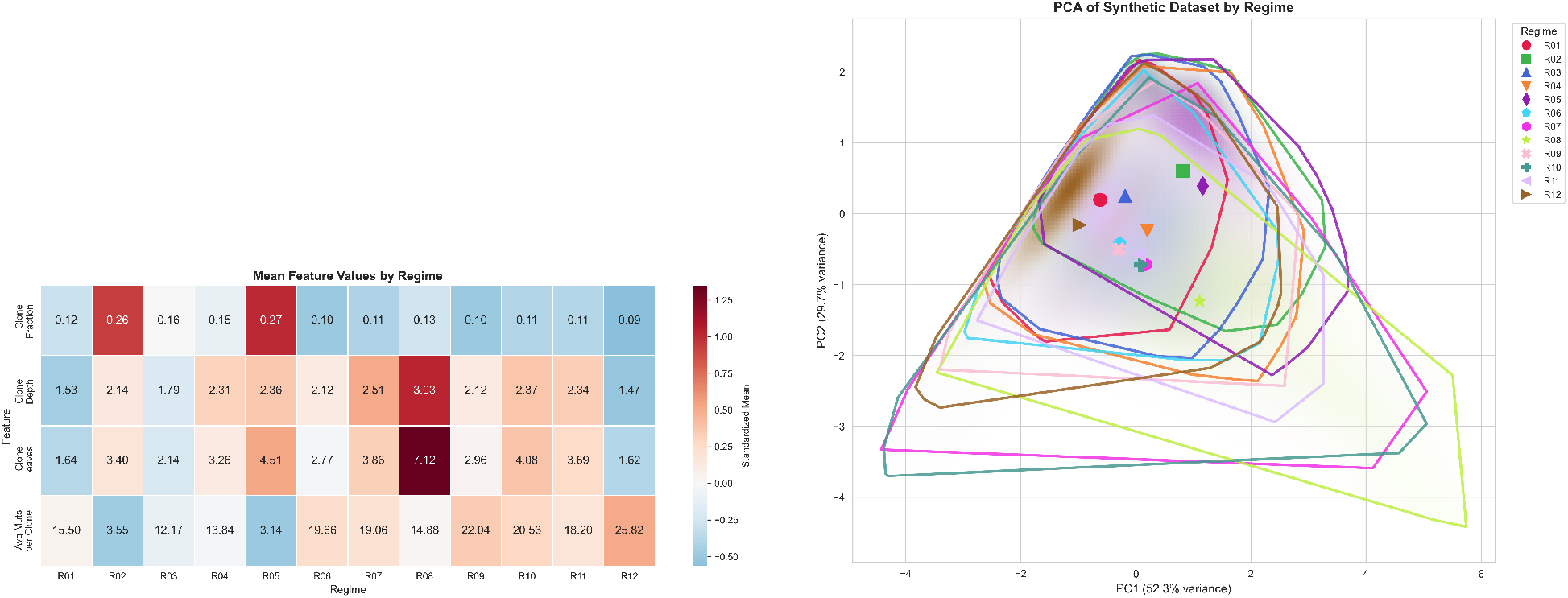
Dataset characterization by simulation regime. (Left) Heatmap of mean feature values— clone fraction, tree depth, clone leaves, and average mutations per clone—across the 12 evolutionary regimes. Regimes R02 and R05 exhibit high clone fractions characteristic of strong selection and chromosomal instability. R08 shows extreme values reflecting complex five-site metastatic dynamics with high migration. R12 (small trees, early detection) has the lowest values across all features. (Right) PCA projection of phylogenetic feature vectors colored by regime. The first two principal components capture the majority of variance, revealing distinct clustering patterns: single-site regimes (R01–R05) occupy one region of the feature space, three-site metastatic regimes (R06–R07) form an intermediate cluster, and five-site metastatic regimes (R08–R10) span a broader region reflecting their greater structural diversity.

## 7 Discussion

### 7.1 Depth Scaling

The 32.1M model (36 layers) diverged across all data fractions, achieving near-zero validity. A likely root cause is optimization at fixed hyperparameters: the learning rate (2 × 10^−4^) and weight decay (10^−12^) were set for the 9-layer model and reused without adjustment. Batch size was also reduced for deeper models (28, 14, 7) to fit in GPU memory, which itself affects optimization dynamics. This was a deliberate choice to isolate model capacity from per-configuration tuning confounders, but scaling depth 4 × without adapting the optimization schedule is known to cause instability — deeper networks have different loss landscape curvature, and learning rates appropriate for shallow networks can cause divergence in deeper ones. Training logs are consistent with this interpretation: the 32.1M model’s validation loss diverged from epoch 0 at 30% and 60% data, and after epoch 149 at 100% data. While it is possible the problem might be resolved by additional training data needed to parameterize the 32.1M model, the pattern of immediate divergence (rather than gradual overfitting) further supports an optimization failure rather than a fundamental architecture–task mismatch, though without successfully training the 32.1M model under adapted hyperparameters, we cannot definitively separate these explanations.

Extending to deeper configurations may likely require learning rate warmup, layer-wise learning rate decay, gradient clipping, and stabilization techniques such as Pre-LayerNorm. None of these were applied: the experiment was designed to isolate raw model capacity for fixed optimization hyperparameters, not to determine the best achievable performance at each scale. Consequently, we cannot distinguish whether the 32.1M failure reflects a fundamental depth–task mismatch or a correctable optimization problem. The 16.2M model’s strong performance suggests that the task may not require extreme depth given our dataset size, but confirming this would require successfully training the 32.1M model under an adapted optimization protocol.

### 7.2 Scaling Regimes

The scaling experiments reveal three capacity regimes with different implications. At the lower end, the 8.2M model learns valid structure reliably (89–94%) but captures less distributional variation, with higher MMD than the 16.2M model; its validity at 100% data (89.7%) is lower than at 60% (94.2%), consistent with the full dataset’s structural diversity exceeding the model’s capacity. The 16.2M model hits a sweet spot, achieving the best balance at 60% data (96.5% validity, lowest MMD), though increasing to 100% data improves validity (97.5%) at the cost of higher MMD, consistent with overfitting to training-specific patterns. At 32.1M parameters, the model fails entirely due to optimization instability rather than inherent overcapacity. These results indicate that for phylogeny generation, the capacity–data–performance relationship is non-monotonic at fixed hyperparameters.

### 7.3 Cross-Regime Transfer

Models trained on diverse regimes learned representations that partially transferred to held-out regimes. The common phylogenetic structure—clone branching, mutation attachment, tree topology—appears learnable from examples across the evolutionary landscape, even when regime-specific details (depth distributions, mutation burdens) do not fully transfer. These transfer effects were observed in a low-data regime (700 graphs); at larger data scales, regime-specific memorization may become feasible, potentially reducing the relative advantage of diverse training.

Single-regime training (R1 Only) achieved the highest in-distribution validity (66.2%) but failed on other regimes, consistent with the model learning regime-specific patterns rather than general validity rules. Metastatic regimes (R06–R10) were harder to generate accurately than single-site regimes, with generation difficulty correlating with structural complexity.

### 7.4 Limitations

Perhaps the most significant limitation of the present work is the simulation-to-real gap: all experiments use synthetic data from the SISTEM simulator to allow for a large and structurally diverse training library. Generalization to patient-derived phylogenies remains challenging to attempt, both in identifying sufficiently large and diverse libraries of patient-derived phylogenies on which to train and in managing novel sources of error arising from such sources as sequence error, phylogeny inference error, and biases in the cases profiled. As a result, the applicability of the strategy here to purely real data remains untested and may face significant domain shift. Semi-synthetic training, augmenting more limited real data with synthetic, may provide one path forward.

Additionally, excluding graphs with more than 200 nodes disproportionately affects metastatic regimes (R08–R11), so results on metastatic structure apply to moderate-complexity trees rather than the full range. More broadly, the *O*(*n*^2^) edge tensor representation limits scalability: extending to phylogenies with several hundred or more nodes would require sparse or hierarchical representations to remain tractable.

A related concern is the coarseness of our evaluation metrics. The four summary statistics (clone fraction, depth, mutations per clone, leaves) may miss fine-grained distributional differences; metrics over full degree distributions, subtree size distributions, or tree edit distances could reveal performance gaps invisible to scalar summaries. We also do not perform a nearest-neighbor analysis between generated and training graphs, which would be needed to distinguish genuine generalization from memorization. Finally, all results are from a single random seed for both data splitting and model initialization, meaning that small differences between configurations (e.g., 1–2 point validity gaps) may not be robust to replication.

Our validity checker may overestimate structural correctness. While it verifies acyclicity, root degree, and edge-type consistency, it does not enforce that each mutation node connects to exactly one clone (degree one), nor does it check full clone-backbone connectivity. Graphs that pass all four checked constraints may still violate these additional properties; reported validity rates should therefore be interpreted as upper bounds. Additionally, the undirected edge encoding discards edge directionality, though direction can be recovered unambiguously by breadth-first search from the designated root node. Finally, the absence of baseline comparisons to other graph generative methods (e.g., autoregressive or flow-based models) limits our ability to contextualize DiPhy’s absolute performance; the results demonstrate feasibility but do not establish relative advantage.

### 7.5 Future Directions

Several directions could extend the model’s capabilities. We deliberately focused here on our ability to create a broadly applicable generative model of tumor phylogeny-like structure, but putting that to practical use will require being able to generate samples conditioned on patient-specific data. Conditioning on observed data (mutation profiles, variant allele frequencies, partial tree structure) would enable prior-guided reconstruction, posterior sampling, and data augmentation conditioned on tumor type. A complementary approach is tree-aware diffusion: noise schedules designed for tree-structured objects (corrupting leaves before roots) or hierarchical denoising that respects parent–child dependencies could improve validity without post-hoc enforcement. Incorporating validity checking directly during the reverse process offers another route to ensuring structural correctness while preserving learned distributional structure. Relatedly, validity-by-construction approaches such as GFlowNet-based samplers that build trees incrementally under hard constraints [24] represent a complementary paradigm; combining learned distributional priors from diffusion with constrained sequential generation could leverage the strengths of both. Combining diffusion-based sampling with more traditional consensus tree methods for tumor phylogenetics [8] could provide another alternative path to generating accurate phylogeny models from imperfect samples. Finally, bridging the gap to clinical relevance will require domain adaptation — pretraining on simulated data followed by fine-tuning on phylogenies inferred from real tumors — to address the simulation-to-real gap identified above.

### 7.6 Conclusion

In the present work, we show that discrete graph diffusion models can be trained to generate structurally valid graphs mimicking tumor phylogenies with low distributional distance to held-out test data. Our experiments show a non-monotonic capacity–performance relationship, a benefit of regime-diverse training for cross-regime transfer in the low-data setting, and partially independent scaling behavior of validity and distributional fidelity. These conclusions are drawn from a single dataset and seed; further validation across additional simulators, real-data benchmarks, and replicated runs will be important to establish their generality.

## Code and Data Availability

Code and data is available at https://github.com/siddsabata/DiPhy.

## AI Usage Acknowledgment

AI-assisted tools (OpenAI GPT, Anthropic Claude, Claude Scientific Skills) were used during the preparation of this manuscript for three purposes: (1) iterating on the articulation and presentation of ideas, (2) generating and debugging code for experiments and analysis, and (3) refining drafts for clarity and structure [13]. All scientific contributions—including study design, hypotheses, interpretation of results, and final conclusions—originated with the authors. All AI-generated or AI-assisted output was critically reviewed, verified, and edited by the authors, who take full responsibility for the content of this work.

## A Simulation Regimes

**Table 4:** Overview of simulation regimes used in this study, showing biological interpretation and planned versus retained sample counts. Regimes span single-site primary tumors and multi-site metastatic settings with varying selection strength, chromosomal instability, migration structure, and detection assumptions.

| Regime | Biological interpretation | Planned / actual |
| --- | --- | --- |
| R01: Single-site, arm-level, near-neutral evolution | Primary single-site tumor dominated by arm-level CNAs with weak or near-neutral selection, approximating drift-like copy-number evolution with few strong sweeps. | 2000 / 1789 |
| R02: Single-site, arm-level, strong selection | Single-site primary tumor with arm-level CNAs under strong selection, producing frequent selective sweeps and later detection at higher tumor burden. | 2000 / 1557 |
| R03: Single-site, region-driven selection | Single-site primary tumor where selection is driven by regional or gene-category composition, yielding heterogeneous CNA fitness effects. | 2000 / 1759 |
| R04: Single-site, hybrid CNA and SNV selection | Single-site tumor with hybrid selection in which both CNAs and SNV drivers contribute to clonal expansions. | 2000 / 1731 |
| R05: Single-site, high chromosomal instability and WGD | Single-site tumor with high chromosomal instability and frequent whole-genome duplication, generating large ploidy shifts and late-stage-like karyotypic chaos. | 2000 / 1396 |
| R06: Three-site metastasis, static anatomy, shared landscape | Three-site metastatic setting with static anatomy and no site-specific fitness landscapes, emphasizing founder effects and low migration. | 1500 / 866 |
| R07: Three-site metastasis, distance-based landscapes | Three-site metastatic setting with distance-based fitness landscapes, introducing site-specific selection structured by anatomical distance. | 1500 / 460 |
| R08: Five-site metastasis, high migration and reseeding | Five-site metastatic setting designed to induce high migration and reseeding, producing complex multi-directional spread and inter-site mixing. | 1500 / 64 |
| R09: Five-site metastasis, genotype-dependent migration | Five-site metastatic setting in which migration depends on genotype compatibility, without explicit organotropism priors. | 1500 / 934 |
| R10: Five-site metastasis, genotype migration with organotropism | Five-site metastatic setting combining genotype-driven migration with distance-based fitness landscapes, producing strong site biases and selective bottlenecks. | 1500 / 394 |
| R11: Seven-site metastasis, sparse sampling | Seven-site metastatic setting producing large, sparse multi-site trees with limited sampling depth per site. | 500 / 134 |
| R12: Small trees, early detection, low sampling | Early-detection, low-burden tumors yielding small trees through very low detectability thresholds and sparse sampling. | 2000 / 1497 |

## B Memory Efficiency

Phylogeny graphs can have up to 200 nodes, requiring 200 × 200 × 3 = 120,000 floats per edge tensor when one-hot encoded. We employ several strategies to manage this memory footprint.

### On-Disk Sharding

Graphs are stored in small on-disk shards rather than a single monolithic file. During training, shards are loaded on demand and cached in memory, reducing startup time and enabling datasets larger than RAM.

### On-the-Fly Encoding

One-hot encoding is performed during batching rather than pre-computed. This trades compute for memory, allowing larger batch sizes by avoiding storage of expanded tensors.

### Marginal Precomputation

Node and edge marginals (*m*_*X*_, *m*_*E*_) are computed once on the training split and stored, preventing information leakage from validation/test splits.

## C Hyperparameter Details

Table 5 provides the complete hyperparameter specification for all experiments.

**Table 5:** Complete hyperparameter specification. All model sizes share identical training hyperparameters except batch size; capacity varies through depth only.

| Category | Parameter | Value |
| --- | --- | --- |
| Architecture | Node hidden dim ($d_X$) | 256 |
| | Edge hidden dim ($d_E$) | 64 |
| | Global hidden dim ($d_y$) | 64 |
|  | Attention heads | 8 |
|  | Layers (8.2M / 16.2M / 32.1M) | 9 / 18 / 36 |
| Diffusion | Timesteps ($T$) | 1000 |
|  | Noise schedule | Cosine |
|  | Transition type | Marginal |
| | Edge loss weight ($\lambda$) | 5 |
| Optimization | Optimizer | AdamW (AMSGrad) |
| | Learning rate | $2 \times 10^{-4}$ |
| | Weight decay | $10^{-12}$ |
|  | Gradient clipping | None |
|  | EMA decay | 0 (disabled) |
| Training | Epochs | 500 |
|  | Batch size (8.2M / 16.2M / 32.1M) | 28 / 14 / 7 |
|  | Validation frequency | Every 50 epochs |
|  | Checkpoint selection | Best validation NLL |
| Data | Train / val / test split | 80% / 10% / 10% |
|  | Split seed | 0 |
|  | Max node count | 200 |
| Evaluation | Generated samples | 1,000 |
|  | Sampling steps | 1,000 (full reverse) |

## D Additional Figures

**Figure 6:**
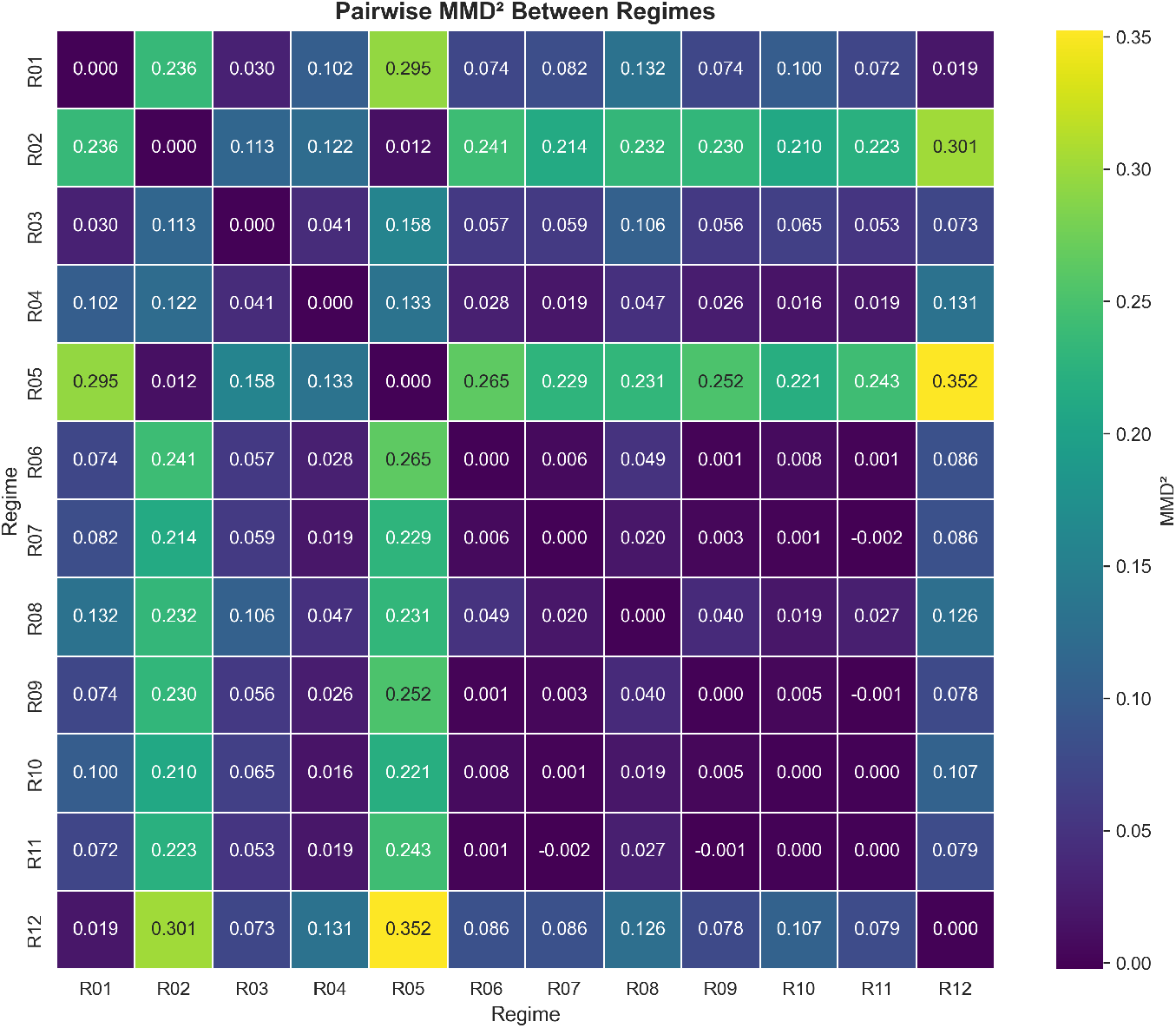
Per-regime distributional similarity for cross-regime models. Heatmap of MMD^2^ between generated samples and test samples stratified by regime, for the three training configurations (Regular, R1 Only, No R1). Darker colors indicate lower MMD (better match). The Regular model shows relatively uniform performance across regimes. The R1 Only model achieves low MMD only on R01, with substantially elevated error on all other regimes. The No R1 model shows low error on trained regimes (R02–R12) but elevated error on the held-out R01, though the transfer gap is smaller than for the R1 Only model’s failure on non-R1 regimes. This asymmetry suggests that diverse training provides more transferable representations than single-regime specialization.

## E Additional Tables

**Table 6:**
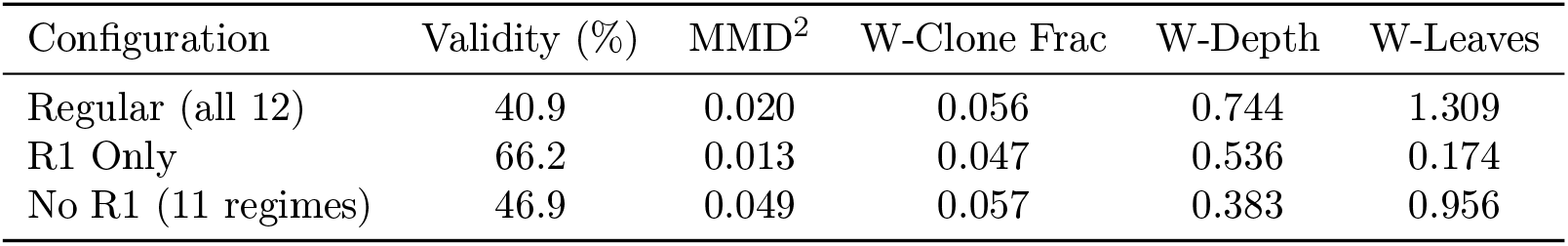
Cross-regime generalization metrics. Summary statistics for the three training configurations evaluated on the full test set across all regimes. Validity is computed on 1,000 generated samples; MMD and Wasserstein metrics compare generated samples to held-out test data.

| Configuration | Validity (%) | MMD<sup>2</sup> | W-Clone Frac | W-Depth | W-Leaves |
| --- | --- | --- | --- | --- | --- |
| Regular (all 12) | 40.9 | 0.020 | 0.056 | 0.744 | 1.309 |
| R1 Only | 66.2 | 0.013 | 0.047 | 0.536 | 0.174 |
| No R1 (11 regimes) | 46.9 | 0.049 | 0.057 | 0.383 | 0.956 |

**Table 7:** Best model configurations by objective. Summary of optimal configurations for different evaluation criteria, highlighting the trade-offs between validity, distributional fidelity, and robustness.

| Objective | Model Size | Data Fraction | Value | Notes |
| --- | --- | --- | --- | --- |
| Highest validity | 16.2M | 100% | 97.5% | Slight increase over 60% data |
| Best MMD | 16.2M | 60% | 0.001 | Optimal model-data balance |
| Most robust | 8.2M | any | 89–94% | Stable across data scales |
| Best W-clone fraction | 16.2M | 60% | 0.006 | Lowest single-feature distance |

## Notes

https://github.com/siddsabata/DiPhy

### Competing Interest Statement

The authors have declared no competing interest.

